# Parametric models for predicting nonstationary spike-spike correlations with local field potentials

**DOI:** 10.1101/2025.07.28.667206

**Authors:** Zeinab Tajik-Mansouri, James P. Dion, Monty A. Escabí, Ian H. Stevenson

## Abstract

**Objective:** Correlations between the spiking of pairs of neurons are often used to study the brain’s representation of sensory or motor variables and neural circuit function and dysfunction. Previous statistical techniques have shown how time-averaged spike-spike correlations can be predicted by the time-averaged relationships between the individual neurons and the local field potential (LFP). However, spiking and LFP are both nonstationary, and spike-spike correlations have nonstationary structure that cannot be accounted for by time-averaged approaches. Here our goal is to develop parametric models that predict spike-spike correlations using a small number of LFP-based predictors and apply these models to the problem of tracking changes in spike-spike correlations over time.

**Approach:** Parametric models allow for flexibility in the choice of which LFP recording channels and frequency bands to use for prediction, and coefficients directly indicate which LFP features drive correlated spiking. Here we demonstrate our methods in simulation and test the models on experimental data from large-scale multi-electrode recordings in the mouse hippocampus and visual cortex.

**Main results:** In single time windows, we find that our parametric models can be as accurate as previous nonparametric approaches, while also being flexible and interpretable. We then demonstrate how parametric models can be applied to describe nonstationary spike-spike correlations measured in sequential time windows. We find that although the patterns of both cortical and hippocampal spike-spike correlations vary over time, these changes are, at least partially, predicted by models that assume a fixed spike-field relationship.

**Significance:** This approach may thus help to better understand how the dynamics of spike-spike correlations are related to functional brain states. Since spike-spike correlations are increasingly used as features for decoding external variables from neural activity, these models may also have the potential to improve the accuracy of adaptive decoders and brain machine interfaces.

## Introduction

Correlations between the spiking activities of pairs of neurons (spike-spike correlations) are key indicators of neural processing (Eggermont, 1990; deCharms and Merzenich, 1996; Salinas and Sejnowski, 2001; Cohen and Kohn, 2011; Panzeri et al., 2022) and dysfunction (Uhlhaas and Singer, 2006). Correlations can indicate similar stimulus-response properties (signal correlations) but can also occur when neurons have shared relationships to intrinsic variables such as local field potentials (LFP). Distinguishing between signal correlations and other shared membrane fluctuations (noise correlations) allows researchers to better predict neural responses and better decode external variables from neural responses (Stevenson et al., 2012). However, one challenge in interpreting neural correlations is that they are typically not fixed but change over time due to changes in the firing rates of the individual neurons and broader changes in stimuli, task, or attentional state (Murthy and Fetz, 1996; Hatsopoulos et al., 1998; Baker et al., 2001; Cohen and Maunsell, 2009; Canolty et al., 2012; Mizuseki and Buzsaki, 2014). Here we aim to develop statistical models to better predict and track time-varying spike-spike correlations by accounting for the relationships between spiking and simultaneously-recorded local field potentials (LFP).

Measuring spike-spike correlations and interpreting their causes can be statistically challenging (Stevenson et al., 2008; Grün, 2009; Gerkin et al., 2013; Harrison et al., 2013). Spike-spike correlations may be due to synaptic connections between the observed neurons and other direct physical links (Moore et al., 1970; Fetz et al., 1991; Swadlow and Gusev, 2001), as well as, indirect effects such as shared synaptic input from other neurons (Swadlow et al., 1998; Trepka et al., 2022) or otherwise correlated input currents (Haider et al., 2016). When measuring correlations in short time windows or when the firing rates of the neurons are low, the number of coincident spikes is small and estimates of correlation have high uncertainty. Additionally, correlations between neurons vary substantially from one pair of neurons to the next – in both the strength of the correlations and their timescales. Here we propose one strategy for modeling these noisy, nonstationary correlations based on parametric models that use features of the LFP as predictors. The relationship between spikes and LFP is complex (Buzsáki et al., 2012; Pesaran et al., 2018; Watson et al., 2018), but broadly speaking, spikes are often coupled to LFPs and occur at specific phases of LFP oscillations. At the same time, spike-spike correlations often have strong oscillatory pattern, and spike-spike correlations appear to be related to the spike-field relationships of the individual neurons (Goldberg et al., 2004; Denker et al., 2011). As LFP spectra change due to brain state, stimuli, and behavior, we expect some of these changes to translate into predictable changes in spike-spike correlations over time.

Many previous studies have used the LFP to predict the activity of single neurons (Harris et al., 2003; Canolty et al., 2010; Kelly et al., 2010), and these studies often find that observed spike-spike correlations are consistent with the model predictions (Zhou et al., 2015; Cui et al., 2016). By modeling the observed correlations directly, previous studies have also developed extensive methods for estimating high-dimensional correlations (Yatsenko et al., 2015; Kass et al., 2023; Nejatbakhsh et al., 2023), separating stimulus and noise correlations (Gawne and Richmond, 1993; Josić et al., 2009; Sadeghi et al., 2019; Keeley et al., 2020), testing for the presence of fast correlations (Amarasingham et al., 2012; Spivak et al., 2022) and putative synapses (Barthó et al., 2004; Kobayashi et al., 2019; Ren et al., 2020), or detecting changes in correlations over time (Lepage et al., 2013). Here we focus on direct, model-based descriptions of the spike-spike correlations that account for the contribution of time-varying LFP. Data analytic tools that can account for the detailed contribution of LFP to spike-spike correlations may be useful in allowing neuroscientists to better understand how and why these correlations change over time and to identify the specific contributions of different anatomical sources and frequency bands. Since LFP-driven correlations are often present alongside stimulus-driven correlations or fast synaptic effects, these data analysis tools may also have the potential to lead to improvements in decoding and more accurate detection of synaptic connections.

Our aim in this work is to predict cross-correlations between spiking neurons based on their individual spike-LFP coupling. Here we first introduce a generalized bilinear modeling framework that can take covariates derived from LFP and predict the observed correlations. We use simulations to illustrate the basic framework and then apply our methods to experimental data from pairs of neurons recorded in mouse hippocampus and cortex. Since the LFP covariates can take many forms and can be based on either single or multiple recording sites, we compare models with different types of covariates and with different optimization strategies. We also compare our parametric models with previously-developed non-parametric methods that predict time-averaged spike-spike correlations based on the spike-triggered average LFP (STA). After applying our methods to time-averaged data, we extend our parametric model framework to the case where both correlations and covariates are calculated over multiple short-time windows. We find that, due to time-varying LFP statistics, the predictions of our models change dynamically over time and partially account for the observed nonstationarity in spike-spike correlations.

## Methods

### Predicting cross-correlograms with LFP features

Given the binned spiking of the two neurons *n*_1_ and *n*_2_ over time, the time-averaged cross-correlogram is given by

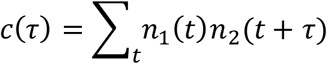

Our goal here is to develop predictions *ĉ*(*τ*) based on observations of the local field potential *y*. In previous work, Halliday et al. (1995) developed an STA Method for constructing predictions based on the concept of the partial correlation or partial spectra. With this approach, we first estimate the spike-triggered average (STA) local field potential for each neuron

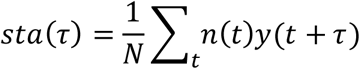

where *N* is number of spikes. The method of partial spectra then constructs the prediction in the frequency domain by taking the cross-spectrum of the STAs for each neuron and whitening it by the spectrum of the LFP *Y*.

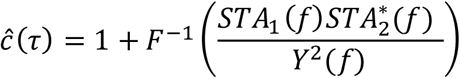

Here *STA*_1_ and *STA*_2_ denote the Fourier transforms of the STAs for the two neurons, and *Y* denotes the Fourier transform of the LFP computed over a common set of frequencies *f*. 𝐹^−1^ denotes the inverse Fourier transform and ⋅^∗^ denotes complex conjugation. Although this model often fits well in practice (Goldberg et al., 2004; Middleton et al., 2012), predictions can sometimes go wrong, especially when the firing rates of the neurons are low or when the LFP power is low at certain frequencies.

Here we consider alternative approaches that take two general forms: an indirect model-based approach and a direct model-based approach. The indirect approach follows the STA Method and fits the correlogram by first describing the rates of the individual neurons *λ*_1_ and *λ*_2_ based on their relationship with the LFP. Then, in a second stage, we use the single-neuron predictions to construct a prediction of the correlation

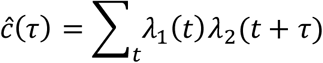

This indirect strategy has been used by several previous studies to understand to what extent rate models of individual neurons and populations of neurons can explain the observed correlations (Cui et al., 2016). In the special case where the rate models are linear, the full prediction can be described in closed form. Suppose we represent the LFP using a set of covariates ***X*** we have *λ*_1_(𝑡) = 𝒙^𝑇^(𝑡)𝜷_1_ and *λ*_2_(𝑡) = 𝒙^𝑇^(𝑡)𝜷_2_, with coefficients given by

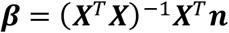

and the correlogram prediction given by

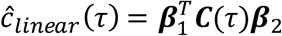

where 𝑪 is a cross-covariance matrix ∑_𝑡_ 𝒙^𝑇^(𝑡)𝒙(𝑡 + *τ*). The LFP covariates ***X*** can take different forms (see below) and can include, for example, band-pass filtered LFP or analytic signals at one or more recording sites. Although the focus of our work here is on LFP, other covariates related to stimulus or task variables could also be included in the set of predictors 𝑋.

In practice, the linear rate estimates *λ* are poor descriptions of nonlinear spiking activity, so some additional modeling is required to make *ĉ* a better match of the observed correlation. In the results here (see below), we consider the specific case where we start with *λ*_𝑖_(𝑡) = 𝒙^𝑇^(𝑡)𝜷_𝑖_, and the correlogram prediction is then modified by an additional output nonlinearity. We use Poisson regression,

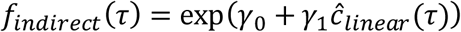

and assume that the number of coincident counts observed in the correlogram is distributed following 𝑐(*τ*)∼𝑃𝑜𝑖𝑠𝑠𝑜*n*(*f*(*τ*)). Our indirect model-based approach thus works by first fitting linear models to the rates with coefficients 𝜷, then, in a second step, fitting an output nonlinearity with coefficients 𝜸 to better match the correlogram. As with the STA method, this approach is based on first describing the relationships between each of the neurons and the LFP. The 𝜷s in our model represent a type of whitened spike-triggered average of the predictors ***X***. However, with this approach we use the time-domain rather than frequency-domain. Additionally, we can compare the impact of different choices for ***X***, and, if needed, use tools from regression and model selection to prevent overfitting.

We also consider a direct approach for cases where we may not need or want descriptions of the individual neurons or their rates over time. In this case, we fit a generalized bilinear model (GBLM)

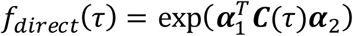

where rather than predicting spike-LFP coupling for each neuron (with 𝜷_1_ and 𝜷_2_), we aim to reconstruct the spike-spike correlation directly by using coefficients 𝜶 = {𝜶_1_, 𝜶_2_}. Here we optimize by coordinate ascent – iteratively fitting 𝜶_1_ with fixed 𝜶_2_then fitting 𝜶_2_with fixed 𝜶_1_ – on the log likelihood, assuming correlogram observations are Poisson distributed with mean *f*_𝑑𝑖𝑟𝑒𝑐𝑡_.

To evaluate the accuracy of model fits we use the pseudo-R^2^

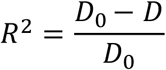

Where 𝐷_⋅_ = 2(ℒ_𝑠_ − ℒ_⋅_) denotes the model deviance for the null (𝐷_0_) and fitted (𝐷) models with log-likelihood ℒ ∝ ∑_*τ*_(𝑐(*τ*) log *f*(*τ*) − *f*(*τ*)). For the null model, *f*(*τ*) is a constant 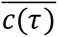, and ℒ_𝑠_ denotes the log-likelihood of the saturated model with *f*(*τ*) = 𝑐(*τ*).

### LFP Representations

One potential advantage of both the indirect and direct modeling approaches described here is that we can choose an appropriate, compact representation of the LFP to use for our predictor 𝑋. Previous work has found that neurons often couple to specific LFP frequency bands and phase-lock to different phases within each band. This observation motivates a generic representation of the form

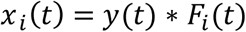

where 𝐹_𝑖_ denotes a bandpass filter applied to the LFP *y*(𝑡), and we choose a set of filters 𝑖 ∈ {1 ⋯ 𝑝} to account for the sensitivity of the two neurons whose spike-spike correlation we aim to model. For modeling single-neuron responses, previous studies have used these types of representations to explain sensitivity to broadband LFP (Truccolo et al., 2005), beta-band rhythms in cortex (Kelly et al., 2010), theta rhythms in hippocampus (Harris et al., 2003), and sensitivity across bands and phases(Cui et al., 2016). Here we consider two models: 1) a compact model based on the Generalized Phase (Davis et al., 2020), and 2) a complex Morlet filterbank model that can capture more-detailed frequency-specific effects.

Briefly, the Generalized Phase (GP) approach (Davis et al., 2020) constructs a broadband analytic signal with a smoothly-varying phase. Recent work (Davis et al., 2022) has shown that the broadband phase is often a better predictor of spiking than the phase of traditional narrowband signals (e.g. theta or beta). Here we consider signals in the band (5-40 Hz, 4^th^-order Butterworth), and the representation takes the form

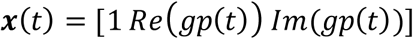

where 𝑔𝑝(𝑡) denotes a complex-valued analytic signal (Davis et al., 2022).

The complex Morelet filterbank has also been previously used for modeling spike-LFP coupling (Cui et al., 2016). Here we use 7 complex-valued Morelet wavelet filters with log-spaced center frequencies varying from 1 to 64 Hz (approximately equal quality factor) plus a real-valued DC. After including an intercept term, this gives a representation ***X*** with 16 predictors.

Note that we can also construct predictors that include features from multiple LFP channels, e.g.

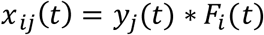

where 𝑥_𝑖𝑗_denotes the output of the 𝑖th filter for the 𝑗th LFP channel. Here we consider GP and filterbank models that use either one LFP channel (the closest channel to neuron 1) or models with two LFP channels (the closest channel to neuron 1 and the closest channel to neuron 2).

### Time-varying prediction

To characterize nonstationary changes in the spike-spike correlation, we measure the correlogramin short-term moving windows. For a time window 𝑤 = (𝑡_𝑠𝑡𝑎𝑟𝑡_,𝑡_𝑒*n*𝑑_), we have observations 𝑐_𝑤_(*τ*) = ∑_𝑡∈𝑤_ *n*_1_(𝑡)*n*_2_(𝑡 + *τ*), and stacking the results from each window generates a data matrix for the overall recording.

Here we are particularly interested in whether modeling LFP allows for dynamic description of the spike-spike correlation. In addition to the global models described above, we thus also consider local observations and models based on calculating and predicting the spike-spike correlation within a moving window. For a time window 𝑤 = (𝑡_𝑠𝑡𝑎𝑟𝑡_, 𝑡_𝑒*n*𝑑_), we have observations 𝑐_𝑤_(*τ*) and predictions *ĉ*_𝑤_(*τ*) or *f*_𝑤_(*τ*) based on the local LFP cross-covariance 𝑪_𝑤_(*τ*) = ∑_𝑡∈𝑤_ 𝒙^𝑇^(𝑡)𝒙(𝑡 + *τ*). Even when the parameters {𝜷, 𝜸} or 𝜶 are fixed, the predictions of both the indirect and direct models described above will change with the spectrum of the LFP.

In general, we find that such stationary models do not accurately track the time-varying correlograms due to the time-varying rates of the two neurons. Rate changes lead to shifts in the baseline of the correlogram that a stationary model cannot track. To better match the data, we consider two methods for tracking these changes. In a first approach, we maintain fixed parameters, but modify predictions with an offset of the form

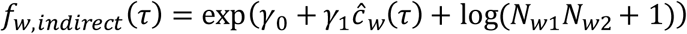

where *N*_𝑤𝑖_ denotes the total number of observed spikes in the window 𝑤, *N*_𝑤𝑖_ = ∑_𝑡∈𝑤_ *n*_𝑖_(𝑡). When 𝜸 = 0 this offset model gives an additively-smoothed description of the expected baseline correlation that varies with 𝑤 but is flat over *τ*. For general 𝛾, the offset acts as a gain on *f*(*τ*) that varies from window to window based on the observed rates of the two neurons.

In a second approach, we consider a fully-adaptive version of the indirect model where the parameters 𝜸 vary along with the moving window. This approach mirrors previous work on Poisson adaptive filtering and smoothing for point-processes (Eden et al., 2004). Briefly, we consider

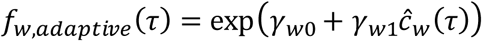

where 𝜸 is updated at each window according to the observed local correlations. Here we track the posterior mean 𝜸_𝑤_ = [𝛾_𝑤0_,𝛾_𝑤1_] and covariance 𝚺_𝑤_ of the parameters using a forward-filtering approach under the assumption that 𝜸 evolves according to 𝜸_𝑤+1_∼*N*(𝜸_𝑤_, 𝝈^𝑻^𝐼). This Poisson adaptive filter has forward updates

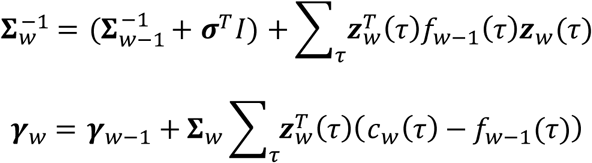

where 𝒛_𝑤_ = [1 *ĉ*_𝑤_]. In this setting, the only additional parameter needed to model the dynamic correlation is the process noise 𝝈, which describes how variable the baseline 𝛾_𝑤0_ and gain 𝛾_𝑤1_ are expected to be from window to window.

### Quantifying stability of cross-correlations over time

To measure the extent of nonstationarity of spike-spike correlations between neuron pairs, we use a low-rank Poisson decomposition and row/column-wise approximations of 𝑐_𝑤_(*τ*). Namely, we fit models of the form

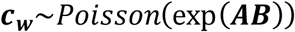

where measured 𝒄_𝒘_ is the [𝑊 × 𝑀] matrix of count observations in 𝑊 non-overlapping windows at 𝑀 delays, with entries 𝑖𝑗 corresponding to 𝑐_𝑤𝑖_(*τ*_𝑗_). The observations are reconstructed based on the combination of a [𝑊 × 𝑘] matrix 𝑨 and a [𝑘 × 𝑀] matrix 𝑩. Here we use 𝑘 ≤ 2 for the low-rank models, and find approximate maximum likelihood estimates of 𝑨 and 𝑩 by coordinate ascent (i.e. alternately optimizing 𝑨 with fixed 𝑩, and optimizing 𝑩 with fixed 𝑨). We also consider the simplified row-approximation exp(𝑎1^𝑇^) where 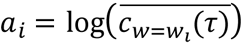 and column-approximation exp(1𝑏^𝑇^) where 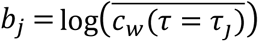. In this case, the predictions are based on the row- and column-wise averages. We then evaluate goodness-of-fit for all models using the pseudo-R^2^ as above.

### Neural Data

To evaluate and test our models, we use multi-region recordings from the Allen Institute for Brain Science’s Visual Coding Neuropixels dataset (https://portal.brain-map.org/explore/circuits). Detailed procedures for surgery, stimulus presentation, and preprocessing are documented in Siegle et al. (2021). Briefly, recordings were obtained from head-fixed, freely-running mice exposed to a variety of visual stimuli, including Gabor patches, full-field drifting gratings, moving dots, and naturalistic images and videos. Our analysis focuses on single-unit data sorted with Kilosort 2, along with simultaneously-recorded local field potentials obtained from the same probe and region as the units. LFP data was low-pass filtered and sampled at 1250 Hz. We restrict our analysis to units with SNR>2 and with more than 1000 spikes. Additionally, we only evaluate recordings with >2hrs of simultaneous spikes and LFP, with electrodes in either the hippocampus or visual cortex, and with ≥10 units per probe in the specified area. Altogether we analyze data from 53 unique recordings, with 44 probes in hippocampus and 51 probes in cortex meeting our criteria. On average, there were 25 units that met criteria per probe in hippocampus and 54 units per probe in cortex across the recordings with 15,000 possible pairs of simultaneously-recorded hippocampal neurons and 86,000 possible pairs of cortex neurons in total. For our population analyses below, we randomly-sampled 25 pairs from each probe with sufficiently-high correlation strength index (CSI) compared to the jittered data, giving *n*=1100 hippocampal pairs and *n* =1400 cortical pairs.

Code for the analysis and simulations described here is available at https://github.com/nztmansouri/nonstationary-correlogram.

## Results

### Predicting spike-spike correlations with LFP

Here we consider models of spike-spike coupling based on simultaneous observations of the local field potential. Intuitively, if the spikes from neuron 1 and 2 have common drive from the LFP, we expect the correlation between neurons 1 and 2 to be partially determined by their separate spike-field relationships. To illustrate how spike-spike correlations can arise from separate spike-field relationships, we first introduce a basic simulation where two Poisson neurons are linearly-coupled to a common local field. Neuron 1 tends to spike at the troughs of the LFP while neuron 2 tends to spike at the peaks (Fig. 1, top). The spike-field relationships are summarized here by the spike-triggered average (STA), and the spike-triggered average is a cross-correlation between the spikes and LFP (De Boer and Kuyper, 1968). At the same time, the two neurons’ separate spike-field relationships give rise to a spike-spike relationship, where the cross-correlation between the two neurons is reduced at delays/intervals near 0, and increased at delays near one-half of the LFP period (Fig. 1, middle).

**Figure 1.**
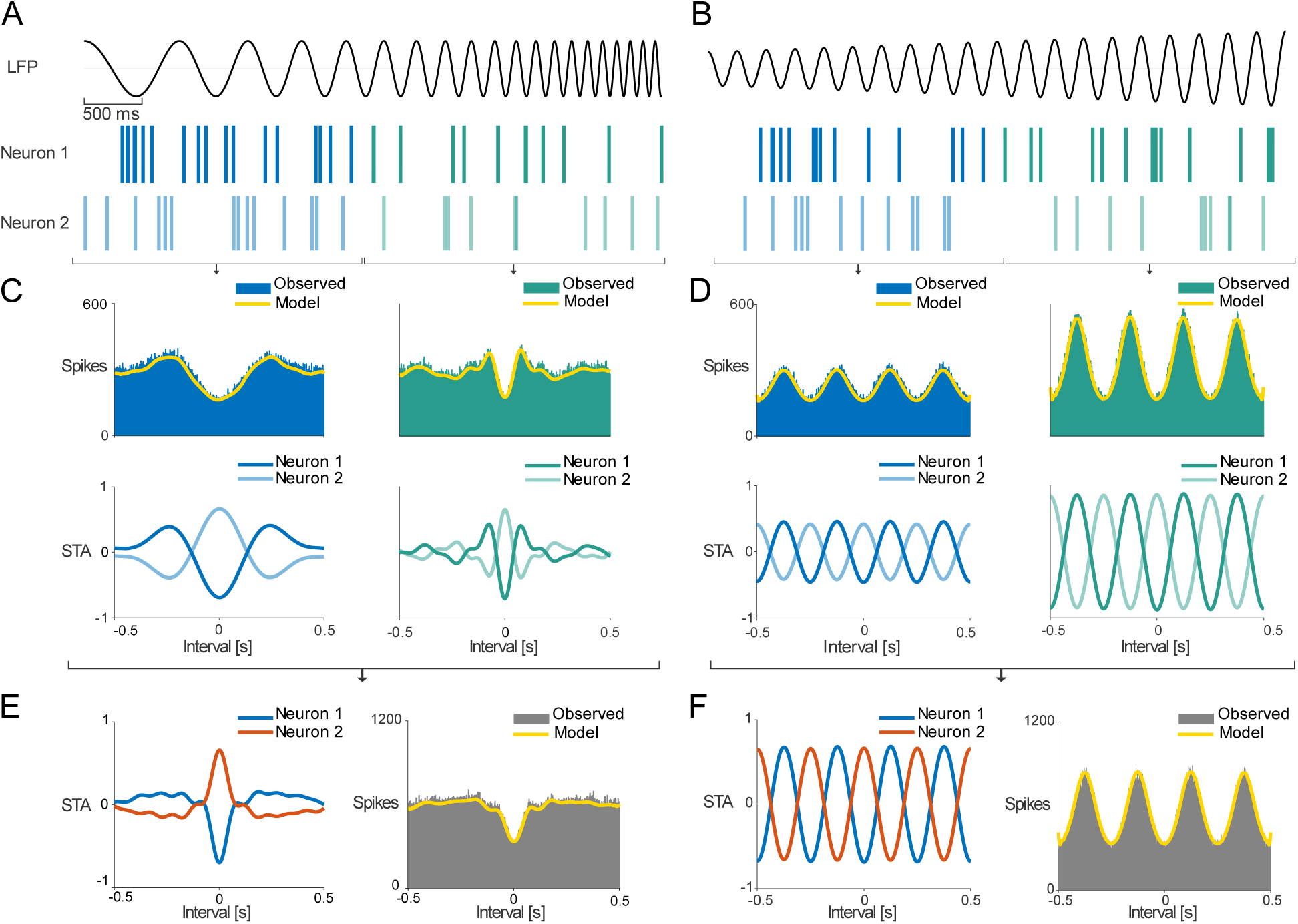
Simulation illustrating how spike-spike correlations can vary with varying LFP. **A)** Simulated LFP (frequency-modulated chirp increasing from 1 Hz to 10 Hz) linearly activates two Poisson neurons, such that neuron 1 spikes at the troughs of the LFP and neuron 2 spikes at the peaks (top). The cross-correlation between the spiking activity of the two neurons (middle) depends on whether we analyze the first half of the data (blue) or the second half (green), and the time-averaged result (bottom) does not necessarily match the results for shorter time windows. A model based on the spike-triggered averages (STA, bottom), accurately captures the spike-spike correlation (yellow). **B)** Same analyses applied to a simulated LFP with constant frequency of 5 Hz, but with modulated amplitude. To reduce noise, the cross-correlations and STAs (in both A and B) are shown with results from 50 trials.

Some analyses have framed the interpretation of spike-spike correlations as one where the observed spike-spike correlation should be distinguished from the partial correlation between the two neurons (Rosenberg et al., 1989, 1998; Eichler et al., 2003). Using the partial correlation rather than the direct pairwise correlation can generate estimates of the excess correlation between the neurons after other variables have been taken into account. However, the partial correlation approach also allows the observed spike-spike correlation to be predicted. When predicting spike-spike cross-correlations at multiple delays given a single LFP, the partial correlation approach works by using the spike-triggered-average LFP (STA) for each of the two neurons. The predicted correlation is then based on the normalized product of the STAs in the frequency domain (see Methods and Fig. 2A). This STA method (Halliday et al., 1995) can often accurately describe the spike-spike correlations observed *in vivo* (Goldberg et al., 2004; Middleton et al., 2012). In Fig. 1, the STA method accurately predicts the spike-spike correlation of the simulated neurons.

**Figure 2.**
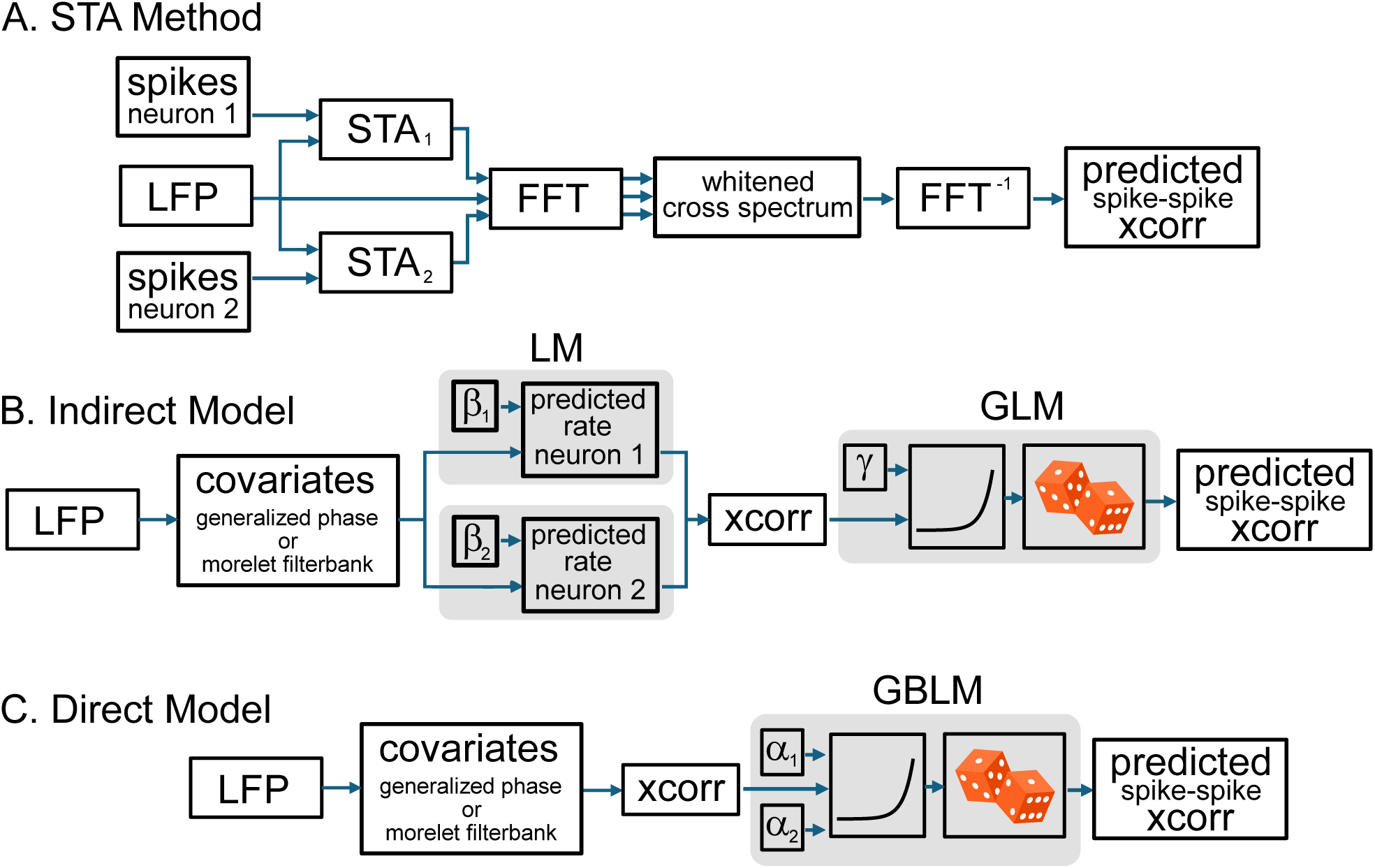
Diagram for the STA method **(A)** and each of the parametric models used here: the indirect method **(B)** and direct method **(C)**.

One potential challenge in predicting spike-spike correlations, in general, is that the LFP may not be stationary over the course of the recording. For a stationary LFP, the STAs and effect of the LFP on the spike-spike correlation will be constant over time as long as the spike-field relationships are fixed. However, for a nonstationary LFP, these quantities can both change with the LFP. Even if the spike-field relationships are fixed, a changing LFP can result in both changing STAs and changing correlations between neurons. In our simulation, we illustrate two potential patterns of changes where the LFP is either modulated in frequency (Fig. 1A) or amplitude (Fig. 1B). In this case, the STA method accurately predicts spike-spike correlations in short windows, but only when the STA is calculated within the same windows. Additionally, although the STA for the time-averaged data (Fig. 1, bottom) can predict the observed time-average spike-spike correlogram, the time-averaged STA does not match the STA for short-windows.The STA method provides a nonparametric approach where data directly defines the predictions. Here we consider alternative parametric approaches for predicting spike-spike correlations and demonstrate how these approaches might be useful for predicting time-varying spike-spike correlations.

To illustrate both the parametric and nonparametric approaches for predicting spike-spike correlations, we consider one example pair of neurons recorded from the hippocampus of an awake mouse along with simultaneously-observed LFP (Fig. 3). Like the simulation, these real neurons are coupled to the LFP and tend to spike at the peaks and troughs of the LFP signal, respectively (Fig. 3A). As with many hippocampal recordings, the LFP signal here has substantial power in the theta band (∼8 Hz), and this oscillation is reflected in the spike-spike correlation. As in the simulation above, the STA method predicts the spike-spike correlation by first measuring the STAs of the two neurons, then normalizing the product of the STAs by the power spectrum of the LFP in the frequency domain (Fig. 3B and Fig. 2A). Similar to previous studies (Goldberg et al., 2004), the STA method accurately predicts the correlation between this particular pair of neurons (Fig. 3C).

**Figure 3.**
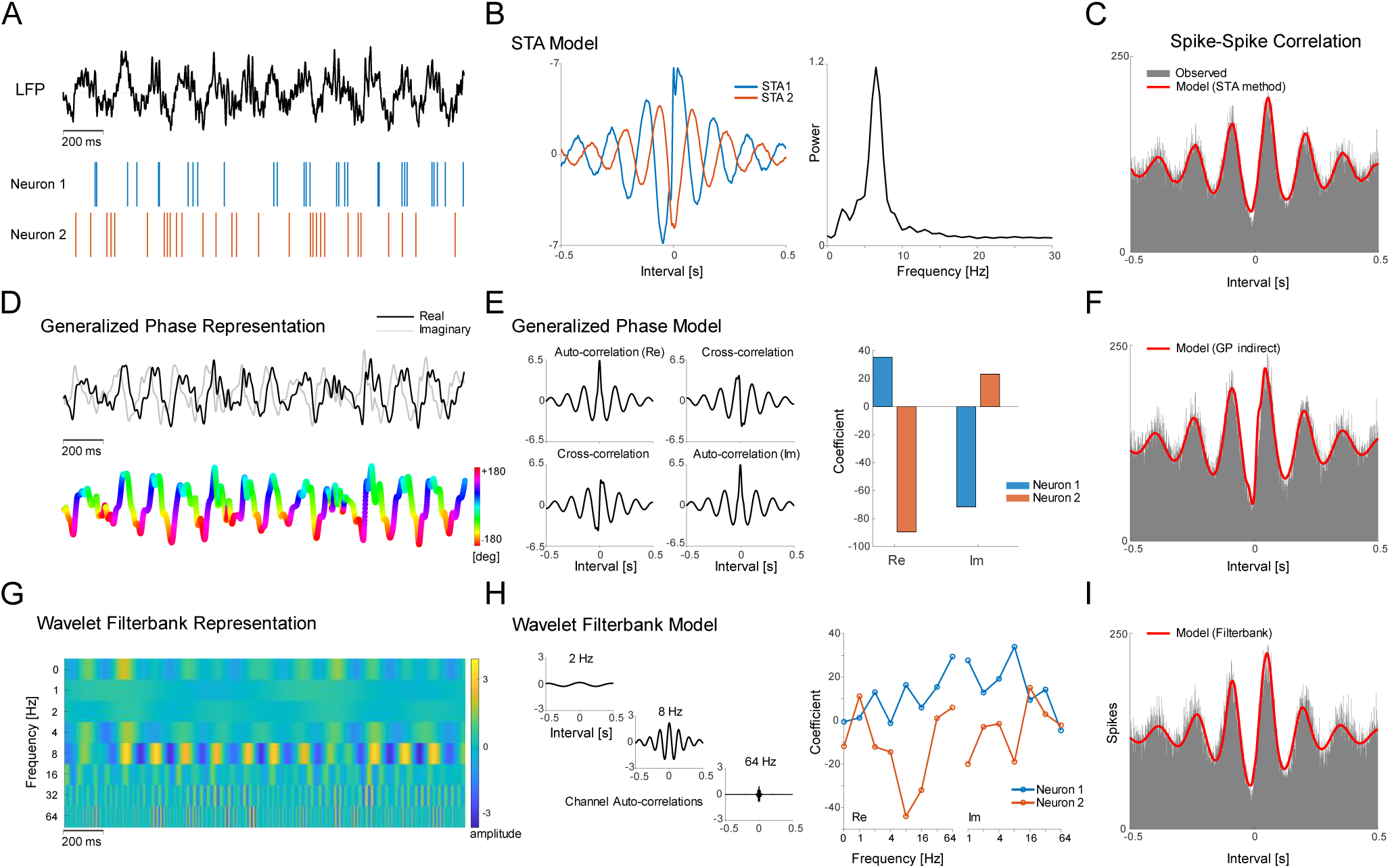
Methods and models for predicting spike-spike correlations from LFP. **A)** An example segment (2 s) of an LFP signal recorded in mouse hippocampus with two simultaneously-recorded neurons coupled to the LFP peaks and troughs. Nonparametric methods for predicting spike-spike correlations are based on the data directly **(B-C)**, while here we consider parametric, bilinear models with an explicit representation of predictors **(D-I)**. **B)** A nonparametric STA method. STAs (left) for neuron 1 (blue) and 2 (orange) are normalized by the LFP power spectrum (right) to generate the predicted spike-spike correlation, with the observed (gray) and predicted (red) correlations shown in **(C)**. **D)** A generalized phase representation where the bandpass-filtered LFP is used to calculate a complex analytic signal ( real part shown in black and imaginary part shown in gray) with smoothly-varying phase (depicted on a color scale below). **E)** The predictions of the parametric model (shown in **F**) are based on the cross-correlation of the predictors (left) weighted by coefficients for each neuron (right). **G)** A wavelet filterbank representation splits the LFP into narrow bands (here the real part is shown), with y-axis indicating the central frequencies of each complex filter. **H)** This representation generates predictors (left) again weighted by coefficients (right) to generate predictions of the spike-spike correlation **(I)**.

As an alternative to the nonparametric STA method, we introduce a parametric approach where an explicit representation of the LFP allows the prediction problem to be framed using a fixed set of predictors weighted by coefficients. Briefly, this approach works by modeling the spike-spike correlations with a bilinear model – where coefficients for both neurons weight the cross-correlation of LFP predictors, followed by a nonlinearity that matches the observed cross-correlation (see Methods and Fig. 2B). To illustrate this parametric approach, we consider two representations for the LFP here: (1) a generalized phase representation (Fig. 3D-F) and (2) a wavelet filterbank representation (Fig. 3G-I). The generalized phase representation extracts two predictors from the LFP, which correspond to the real and imaginary parts of the broadband analytic signal. This model only has two coefficients for each neuron (Fig. 3E), but combiningthese coefficients with the cross-correlationof the predictors can yield predictions of the spike-spike correlation nearly as accurate as those of the nonparametric STA method. Similarly, we can consider separating the LFP into narrower frequency bands (Fig. 3G). Here we use a wavelet filterbank model with seven narrow-band, complex Morlet wavelet filters (center frequencies starting from 1 Hz and increasing to 64 Hz, log-spaced octave wide filters) to generate predictors. This model has 15 fixed parameters for each neuron (Fig. 3H), and predicts the spike-spike correlation accurately as well (Fig. 3I). The differences between the coefficients of the two neurons reflect the difference in the neurons’ spike-field relationships, including their amplitude, phase, and sensitivity to specific frequency bands.

There are several approaches to optimize the coefficients of the parametric models. Here we consider an indirect method and a direct method (see Methods). In the indirect method (Fig. 2B), the per-neuron coefficients and the nonlinearity are optimized sequentially, with the per-neuron coefficients fit based on linear prediction of the spikes of each neuron. In the direct method, the coefficients of a generalized bilinear model are optimized directly to predict the observed cross-correlation (Fig. 2C). Results for the indirect method are shown in Fig. 3. In both approaches, the key feature of the parametric models is that they explicitly distinguish between LFP-based predictors and parameters that describe the spike-field or correlation-field relationships. This distinction allows us to directly extend the models to include multiple LFP recording channels, and to make predictions of spike-spike correlations for arbitrary LFP without needing to track or update the STA based on newly-observed spikes.

### Goodness of Fit for Experimental Data

Although both the nonparametric STA method and the parametric models accurately fit this example pair of neurons, the models do not always explain the variability in the data for different pairs of neurons. One challenge in measuring goodness-of-fit for these models is that different pairs of neurons have correlations of different strengths (Fig. 4A-B), and for weak correlations the variance of the cross-correlation at different intervals is mostly driven by noise/chance. To quantify the correlation strength, we use a correlation strength index (CSI) based on the ratio of the variance and the mean of the spike-spike cross-correlations over different delays. Since independent Poisson neurons will have cross-correlations with a variance equal to the mean, a CSI of 1 represents a case where we would not expect the model to predict any meaningful structure in the cross-correlation. Here we measure the CSI of many simultaneously-recorded pairs of neurons in mouse hippocampus (n=1100 pairs) and cortex (n=1400 pairs) from the Allen Institute’s Visual Coding Neuropixels dataset. We find a wide range of CSIs with strong, highly-structured cross-correlations (Fig. 4A-B, top) as well as weaker cross-correlations with no clear variation across different delays (Fig. 4A-B, bottom). The distribution of CSIs is highly-skewed with a median of 2.90 for hippocampus and 2.65 for cortex (Fig. 4C-D). For comparison, we also evaluate the CSI for a null model of each pair of neurons where we jitter the neural spike times (normally-distributed jitter with 1s s.d.). Since neuron pairs with low CSIs (similar to the CSI for the jittered data) do not have substantial structure for the LFP-based models to capture, we focus our analysis on the pairs with CSIs higher than the decision boundary between the jittered and observed CSI distributions. We fit our models to a random sample of 25 pairs from each recording probe with n=1100 pairs from hippocampus and n=1400 pairs from cortex. As expected, we find that the goodness-of-fit (pseudo-R^2^) is correlated with the CSI (Fig. 4C-D).

**Figure 4.**
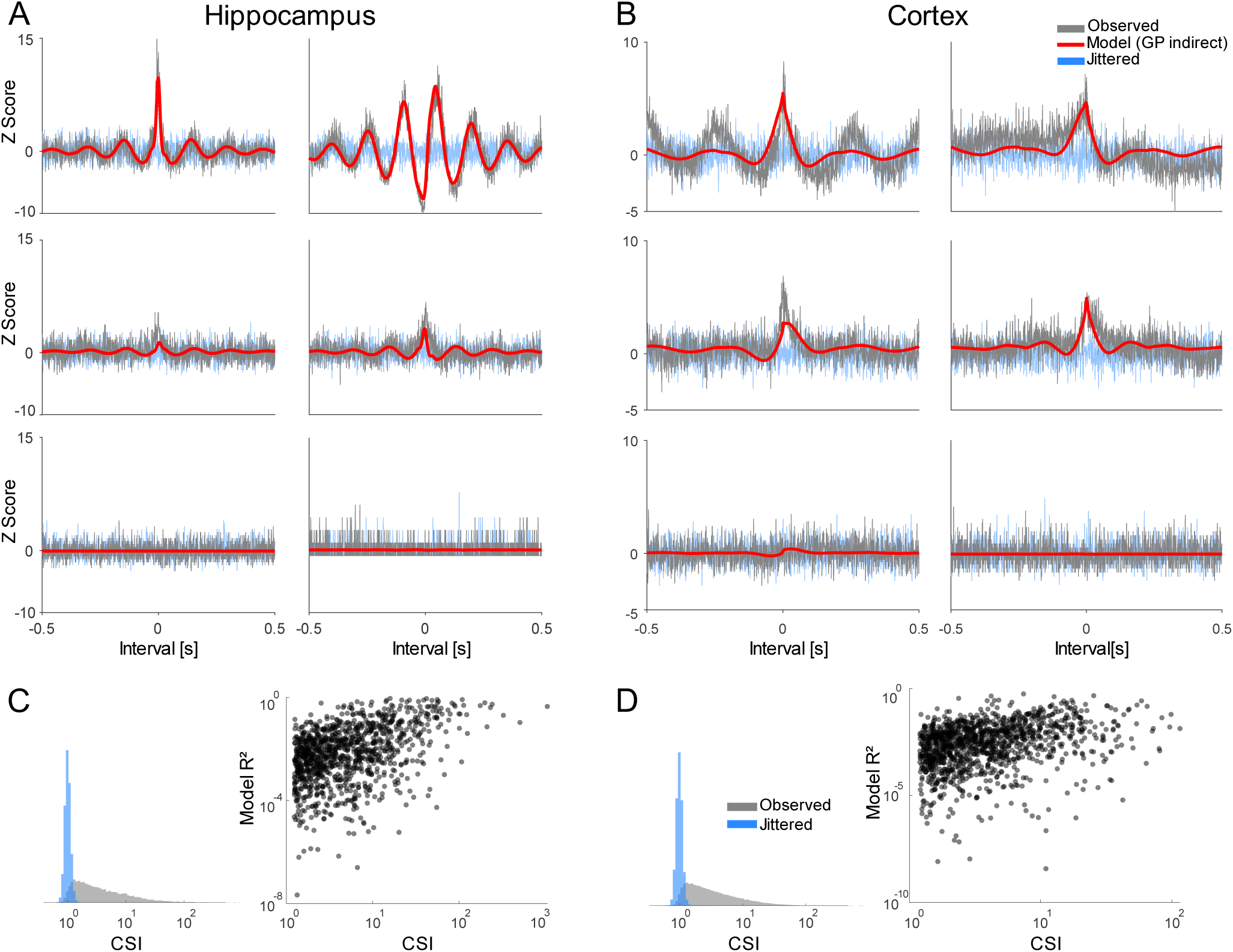
Example cross-correlograms for neuron pairs from the hippocampus **(A)** and cortex **(B)**. Observed correlations (gray) are shown normalized by the correlations of a jittered null model ( blue). The prediction of the indirect model with generalized phase representation of the LFP is shown in red. Histograms of observed correlation strength index (CSI) for all correlogram samples are shown in gray alongside jittered correlograms (blue) for neuron pairs in the hippocampus **(C)** and cortex **(D**). Scatter plots show the model pseudo-R² with respect to CSI for a sample of neuron pairs with CSIs above that of the jittered data.

Using this sample of high-CSI pairs, we compare the performance of the nonparametric STA method to that of several parametric models: 1) a generalized phase (GP) model based on the LFP channel closest to one neuron in the pair, 2) a GP model based on two LFP channels – the closest to neuron 1 and the closest to neuron 2, 3) a wavelet filterbank model of one LFP channel, and 4) a GP model based on one LFP channel fit by direct optimization of the bilinear coefficients. These comparisons vary in terms of the representation of LFP predictors (GP vs filterbank), the number of LFP channels used, and the method of optimizing the coefficients (indirect vs direct). They also differ in the number of parameters (the 1-channel GP indirect model has 8 parameters, the 2-channel GP indirect model has 12, the 1-channel filterbank model has 32, and the 1-channel GP direct model has 6).

Fitting each of these models to the high-CSI pairs from hippocampus and cortex (Fig. 5A and B), we find that the models are all correlated in their goodness-of-fits (pseudo-R^2^), with the results depending on the CSI of the data. However, there are some differences in performance across models. For hippocampus, the 1-channel GP direct model (pseudo-R^2^=0.022 [0.008, 0.069], median and inter-quartile range), 1-channel filterbank model (pseudo-R^2^=0.015 [0.003, 0.068]), and STA method (pseudo-R^2^=0.014 [0.003, 0.070]) all perform well, while the 1-channel (pseudo-R^2^=0.008 [0.002, 0.040]) and 2-channel (pseudo-R^2^=0.010 [0.002, 0.048]) GP indirect models perform slightly worse (Fig. 5A). For cortex, the 1-channel filterbank model (pseudo-R^2^=0.026 [0.005, 0.085]), and STA method (pseudo-R^2^=0.027 [0.006, 0.088]) perform best, while the 1-channel (pseudo-R^2^=0.004 [0.001, 0.012]) and 2-channel (pseudo-R^2^=0.003 [0.001, 0.012]) GP indirect and GP direct (pseudo-R^2^=0.011 [0.005, 0.026]) models perform slightly worse (Fig. 5B). The absolute R^2^ values may be somewhat misleading since the goodness-of-fit is highly dependent on the CSI (Fig. 5, insets). The models, overall, can accurately capture structure in the cross-correlations when there is structure present (e.g. high-CSI examples in Fig. 4).

**Figure 5.**
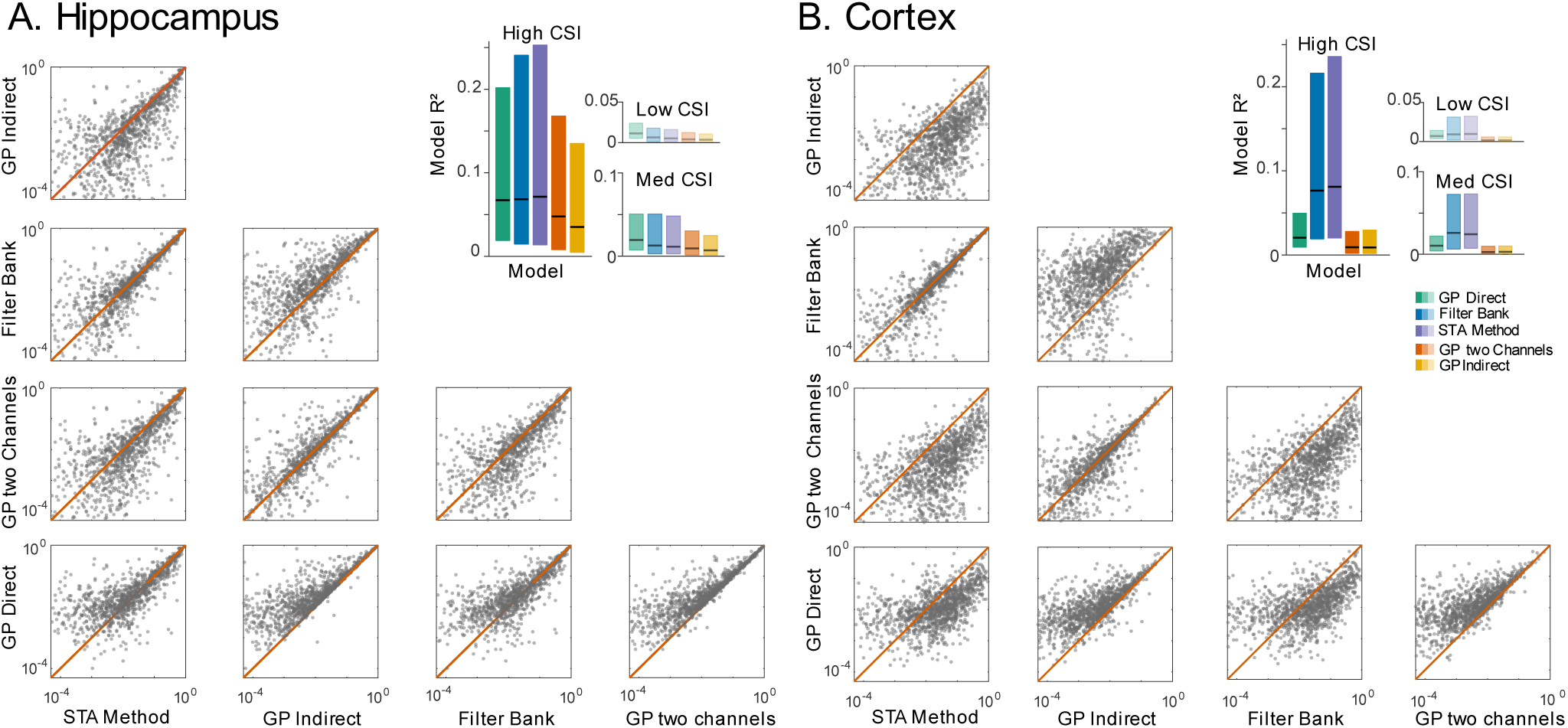
Goodness-of-fit for five models of spike-spike correlations based on LFP: STA method, 1-channel generalized phase model with indirect optimization of the coefficients, 1-channel filterbank model with indirect optimization of the coefficients, 2-channel generalized phase model with indirect optimization of the coefficients, and 1-channel generalized phase model with direct optimization of the coefficients. Scatter plots comparing pseudo-R^2^ for different pairs of models for **A)** hippocampus and **B)** cortex. Insets show dependence of pseudo-R^2^ on data CSI. Boxplots compare the model pseudo-R^2^ measured across different CSI ranges. High, Med, and Low CSI denote the first, second, and third CSI terciles, respectively. Boxplot horizontal black lines denote the median pseudo-R^2^, and the box ranges indicate the interquartile ranges for goodness-of-fits.

For detailed comparison we include the mean pseudo-R^2^ for each CSI level in Table 1. Another consideration in comparing these different models is the computational cost. We calculated average runtime per pair for a sample of neuron pairs on an Intel Core i7-4770 CPU (3.40GHz) with 16GB RAM. These runtimes also depend directly on the recording length (129 for HC and 161 for CTX here) and sampling rate (1250 Hz here). The parametric models are, generally, about an order of magnitude slower than the STA method. Note, however, that this runtime includes time filtering the LFP or calculating generalized phase and, for the indirect models, time for fitting the linear response of each neuron. Since these computations can be shared across pairs that include the same neurons or use the same LFP covariates, there may be substantial room for optimization. Nonetheless, in time-sensitive applications the STA method may still be preferable to our parametric models.

**Table 1:** R2 goodness-of-fit for each model and terciles of correlation strength index, as well as average runtime per-pair.

| Models | Hippocampus |  |  |  |  | Cortex |  |  |  |
| --- | --- | --- | --- | --- | --- | --- | --- | --- | --- |
|  | High CSI | Med CSI | Low CSI | Runtime (s) |  | High CSI | Med CSI | Low CSI | Runtime (s) |
| GP Direct | 0.067 | **0.019** | **0.011** | 5.8 |  | 0.020 | 0.010 | 0.007 | 5.5 |
| Filter Bank | 0.068 | 0.013 | 0.006 | 10.4 |  | 0.076 | **0.026** | 0.009 | 10.1 |
| STA method | **0.071** | 0.011 | 0.005 | 0.6 |  | **0.081** | 0.024 | **0.010** | 0.4 |
| GP two channels | 0.048 | 0.009 | 0.004 | 12.3 |  | 0.009 | 0.002 | 0.002 | 11.7 |
| GP Indirect | 0.035 | 0.007 | 0.003 | 6.3 |  | 0.009 | 0.003 | 0.001 | 5.6 |

### Quantifying non-stationary spike-spike correlations

As in the simulation, a major consideration when predicting spike-spike correlations from LFP is that the LFP is not stationary. LFP power spectra change over time, and we expect these changes to be reflected in changing spike-spike correlations. To determine the nonstationarity of spike-spike correlations, we first measure correlograms for the high-CSI neuron pairs in short time windows of 60s (Fig. 6A-B). Most pairs show dynamic patterns where the correlation increases and decreases over time, and the correlations at different intervals/delays are not fixed over the course of the recording (Fig. 6A-B). Here we are measuring the “unnormalized” correlogram based on spike counts and without adjusting for changes in firing rates over time. When the firing rates of the two neurons vary over the course of the recording, the cross-correlation naturally changes in magnitude. Such changes are not strictly due to changes in LFP, since they can occur even if the neurons are statistically independent (e.g. two independent inhomogeneous Poisson processes).

**Figure 6.**
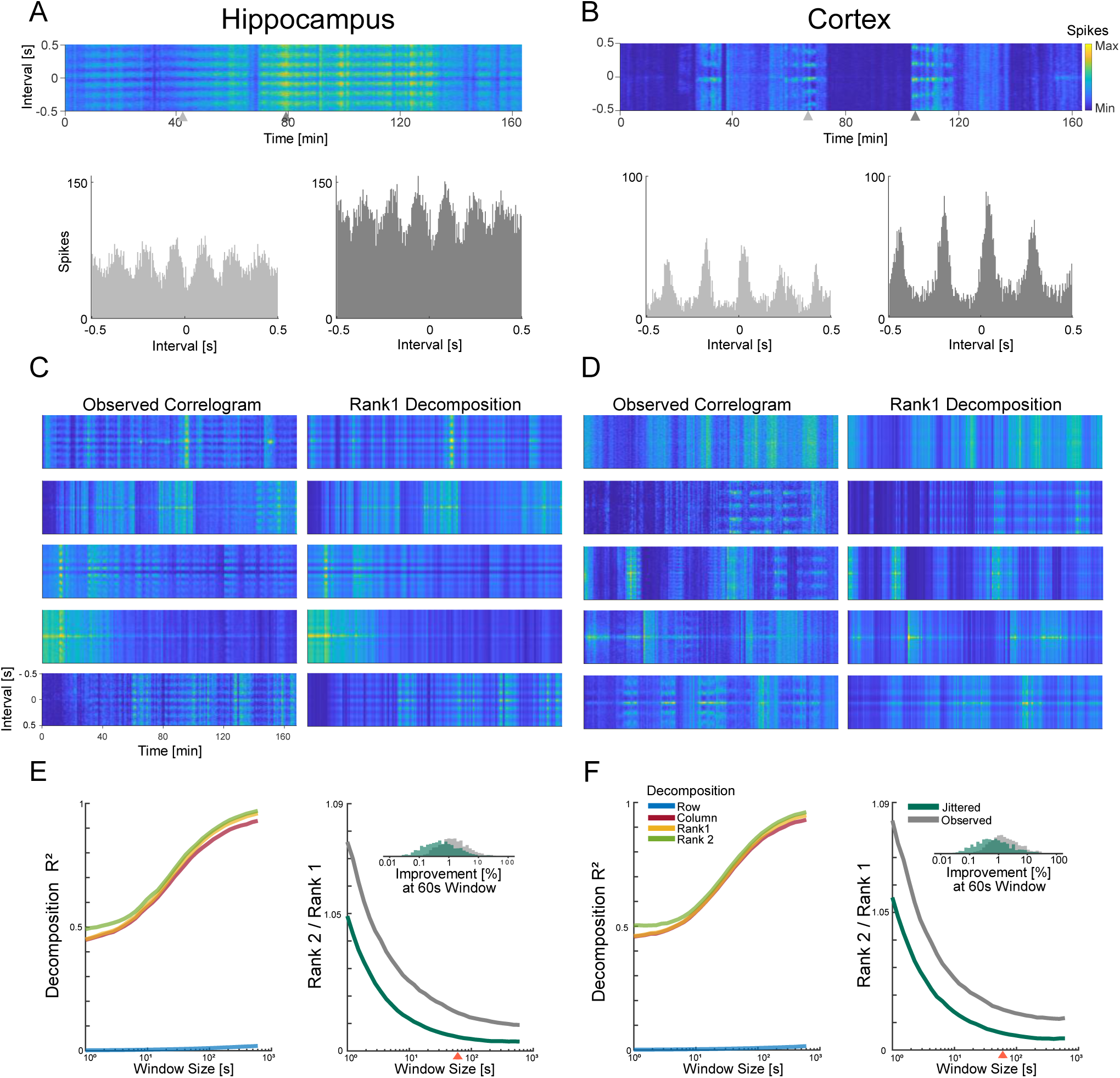
Measuring nonstationarity in spike-spike correlations with matrix decomposition. To illustrate how correlograms change over the course of the recording, we calculate correlations in short time windows. Here, heatmaps indicate correlations at different delays (y-axis) calculated in 60-s time windows over the course of the ∼3hr recording (x-axis). An example of time-varying correlograms are shown for the hippocampus **(A)** and cortex **(B)**, along with correlograms from two single time windows (time indicated by gray triangles). **(C and D)** show additional examples of observed correlogram heatmaps along with a low-rank (rank=1) matrix decomposition (**C**, right for hippocampus, and **D**, right for cortex). For hippocampus (**E**, left) and cortex (**F**, left), pseudo-R^2^ is shown for the row (blue), column (red), rank-1 (yellow), and rank-2 (green) decompositions as a function of window size.(**E**, right and **F**, right) Improvement of the decomposition when using rank-2 compared to rank-1 with both the jittered (green) and the observed correlations (gray). The red arrow indicates a window size of ∼60s, which corresponds to the inset histogram.

To test whether the observed changes in the correlograms over the course of the recording go beyond simple rate changes, we here fit four Poisson matrix decompositions (see Methods). Specifically, we fit: 1) a row-decomposition that assumes an interval-dependent correlation that is fixed over time, 2) a column-decomposition that assumes a time-dependent correlation that is constant for all intervals, 3) a rank-1 decomposition that assumes the matrix can be decomposed as a product of one pattern over time and one over interval, and 4) a rank-2 decomposition. We measure pseudo-R^2^ for each of these matrix decompositions as a function of the window size (Fig. 6E-F). We find that the row decomposition does not accurately track the unnormalized correlograms, while the other three decompositions do well across different window sizes. Most of the variability in these matrices is explained by a column effect where rate fluctuations drive changes in the baseline cross-correlation over the course of the recording. However, a rank-1 decomposition does slightly better by assuming one pattern of correlation across different intervals, and a second pattern that acts as a gain for different time windows (Fig. 6C-D). These decompositions are consistent with the idea that there are fixed patterns of correlations that change due to the changing firing rates of the two neurons. However, we find that a rank-2 decomposition does lead to small improvements in the reconstruction relative to the rank-1 decomposition, suggesting that there may be some additional nonstationarity beyond what results from changing firing rates (Fig. 6E-F).

The goodness-of-fit increases with the window size for all decompositions (Fig. 6E-F, left). For short time windows (< ∼10s) the pseudo-R^2^ for all reconstructions is relatively low, likely due to the fact that the CSI is also lower at this resolution. The improvement provided by the rank-2 decomposition compared to the rank-1 decomposition gives an approximate lower-bound for how much the correlation pattern changes over the course of the recording(Fig. 6E-F, right). For comparison, we also evaluate the improvement from using the rank-2 vs rank-1 decomposition for jittered data (Gaussian noise used to jitter spike times with 1s s.d.). The improvement from using the rank-2 vs rank-1 decomposition for the observed data is higher than the improvement with the jittered data for all window sizes. For 60-s windows (Fig. 6E-F, inset), the rank-2 decomposition shows an improvement over the rank-1 decomposition of 1.4% (median) and 1.3% for cortex and hippocampus, respectively, and the improvement is higher than that of the jittered data (0.6% and 0.5% for cortex and hippocampus). As with the case of predicting stationary spike-spike correlations, CSI is also an important consideration when evaluating nonstationary correlations. This analysis suggests that in some cases there is nonstationarity in spike-spike correlations, and thus our goal here is to test whether these changes can be predicted by the LFP.

### Modeling non-stationary spike-spike correlations with LFP

To illustrate how changes in the LFP spectrum over time might lead to changes in the spike-spike correlations, here we consider one example pair of neurons from the hippocampus (Fig. 7). As the theta-band power and peak frequency vary over the course of the recording (Fig. 7A), the spike-spike cross-correlation for this pair of neurons tends to vary as well (Fig. 7A, middle). We fit our indirect generalized phase model using this LFP channel and find that when we apply the predictions in short 60-s windows (see Methods, Fig. 7A, bottom), the model accurately tracks changes in the correlogram, whichcorrespond to changes in both the LFP’s peak frequency (Fig. 7B) and amplitude (Fig. 7C).

**Figure 7.**
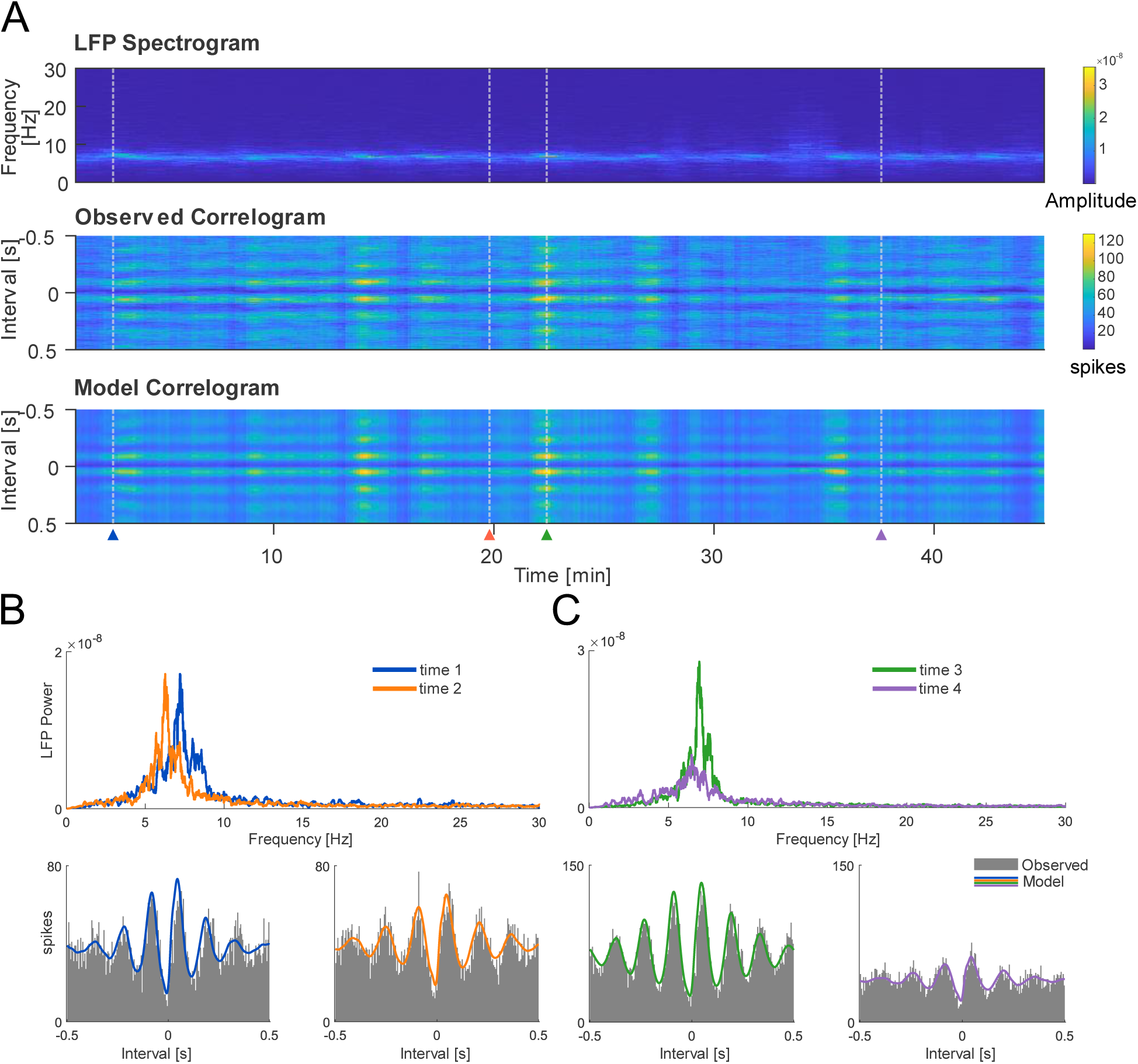
Example and model fits illustrating how nonstationary spike-spike correlations are related to changes in the LFP spectra, with LFP and neuron pair from the hippocampus. **(A)** Top, spectrogram of the LFP signal. Middle, observed correlograms calculated in 60s short-time windows (50-s overlap). Bottom, time-varying correlogram predictions from the indirect generalized phased model. **(B)** Top, the LFP’s PSD for two time points (1=blue and 2=orange) indicating that the dominant frequency of time 1 is higher than that of time 2. In the bottom left and right, the observed correlograms (gray) and corresponding model fits in blue (bottom left) and orange (bottom right) also illustrate the changes in frequency (i.e. by counting the number of peaks if we consider the fluctuations as pseudo-sinusoid). **(C)** Top, the LFP’s PSD for two different time points, in which time 3 (green) has a higher amplitude than time 4 (purple). In the bottom left, the observed correlogram (gray) has higher amplitude (model fit in green) than the observed correlogram (gray) in the bottom right (model fit in purple). Both fits use the generalized phase model.

As with modeling the stationary correlograms, there are multiple possible approaches when describingthe time-varying correlograms that differ in predictors (i.e. generalized phase vs filterbank), optimization strategies (i.e. indirect vs direct) and number of parameters. One major challenge, however, is in how to account for the time-varying rates of the two neurons and the column-wise structure of the matrices described above. Here, when fitting the time-varying indirect model, we assume that there is a known count offset for each window (see Methods). This approach may be appropriate for offline modeling, but a different strategy may be needed when correlations are predicted in real-time. To illustrate one possible approach, we consider an adaptive filtering model to track nonstationary spike-spike correlations from streaming data.

Adaptive filtering has been previously applied to the estimation of changing neural tuning properties (Eden et al., 2004) and synaptic connections (Stevenson and Kording, 2011) from spike data. Here we consider the adaptive filtering approach for LFP-driven spike-spike correlations. Briefly, we use the predictors from the indirect model and add an output nonlinearity with time-varying parameters for the baseline correlation and gain (see Methods). We maintain a distribution over parameters at each time window, and update the mean and covariance as new observations come in. To validate this approach, here we use experimentally-observed hippocampal LFP to simulate two neurons with a spike-spike correlogram with known time-varying gain (Fig. 8). The adaptive filtering model appropriately accumulates uncertainty even when spike observations are missing, and makes predictions based on the LFP. When there are true changes in the baseline or gain, the adaptive filter can track these changes, and distinguish these effects from changes in the LFP itself. Unlike the indirect model described above, the adaptive filter does not require a separate offset when accounting for changes in the column-wise structure. Instead, the column-wise structure is captured by a baseline that is updated based on new observations as they occur.

**Figure 8.**
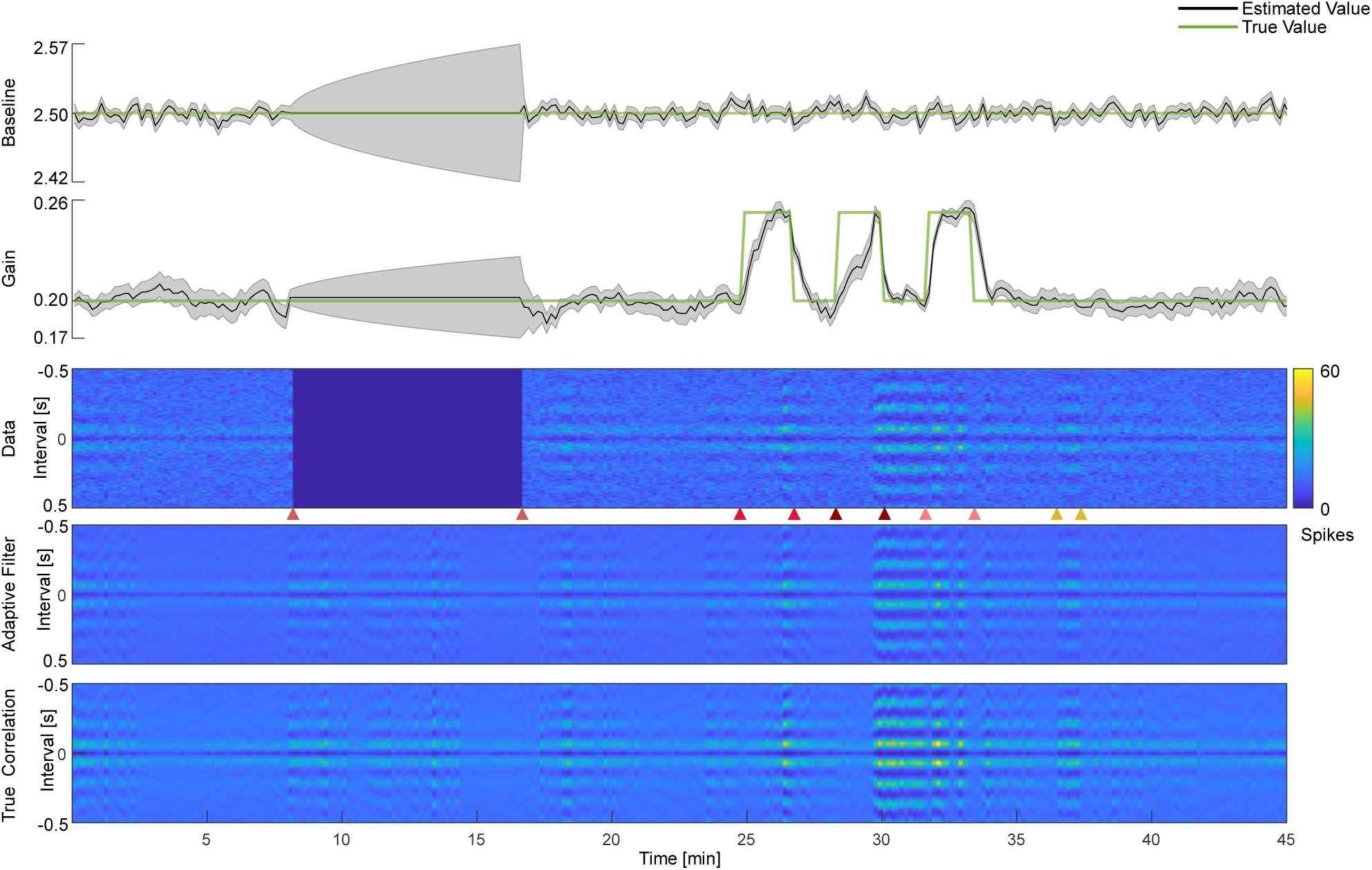
Adaptive filtering model for time-varying correlogram data. Here we use the observed LFP to simulate time-varying spike-spike correlations with known changes in the parameters. We simulate Poisson spiking with a fixed baseline and variable gain (in green in top two rows, respectively) as well as some missing data (time points 8-17 min). The adaptive filter tracks the observed spiking (middle row) over time and updates parameter estimates based on the observations (top two rows, gray bands). The adaptive filter can fill-in missing data (red arrows at time points 8 and 17 min.), and accurately tracks imposed changes on the gain (stepwise changes indicated by arrows at time points between 20 and 40 min). The correlations estimated by the adaptive filter (row above bottom) follow the true correlations (bottom row) when the ground truth is known. Note that some changes in the data are driven by changes in the LFP (e.g. yellow arrow) rather than changes in the adaptive parameters.

To evaluate the goodness-of-fit for these nonstationary LFP-driven models of spike-spike correlations, we again calculate pseudo-R^2^ after fitting the high-CSI pairs from hippocampus and cortex. We compare the nonstationary, indirect generalized phase model and the adaptive filtering model against a stationary model where the same LFP-based prediction (indirect GP) is applied to all time windows. Generally, we find that the nonstationary models are substantially more accurate than the stationary model, and the indirect approach with column-wise rate offsets performs slightly better than an adaptive filtering approach with running parameter estimates (Fig. 9). Using 60-s windows and hippocampal neuron pairs, we obtained median pseudo-R^2^ for the stationary (0.001), nonstationary indirect (0.45), and adaptive filtering models (0.35). For cortical neuron pairs, we also observed median pseudo-R^2^ for the stationary (0.0001), nonstationary indirect (0.41), and adaptive filtering models (0.31). Thus, for both hippocampal and cortical neural pairs, the LFP can provide an accurate prediction of time-varying cross-correlations.

**Figure 9.**
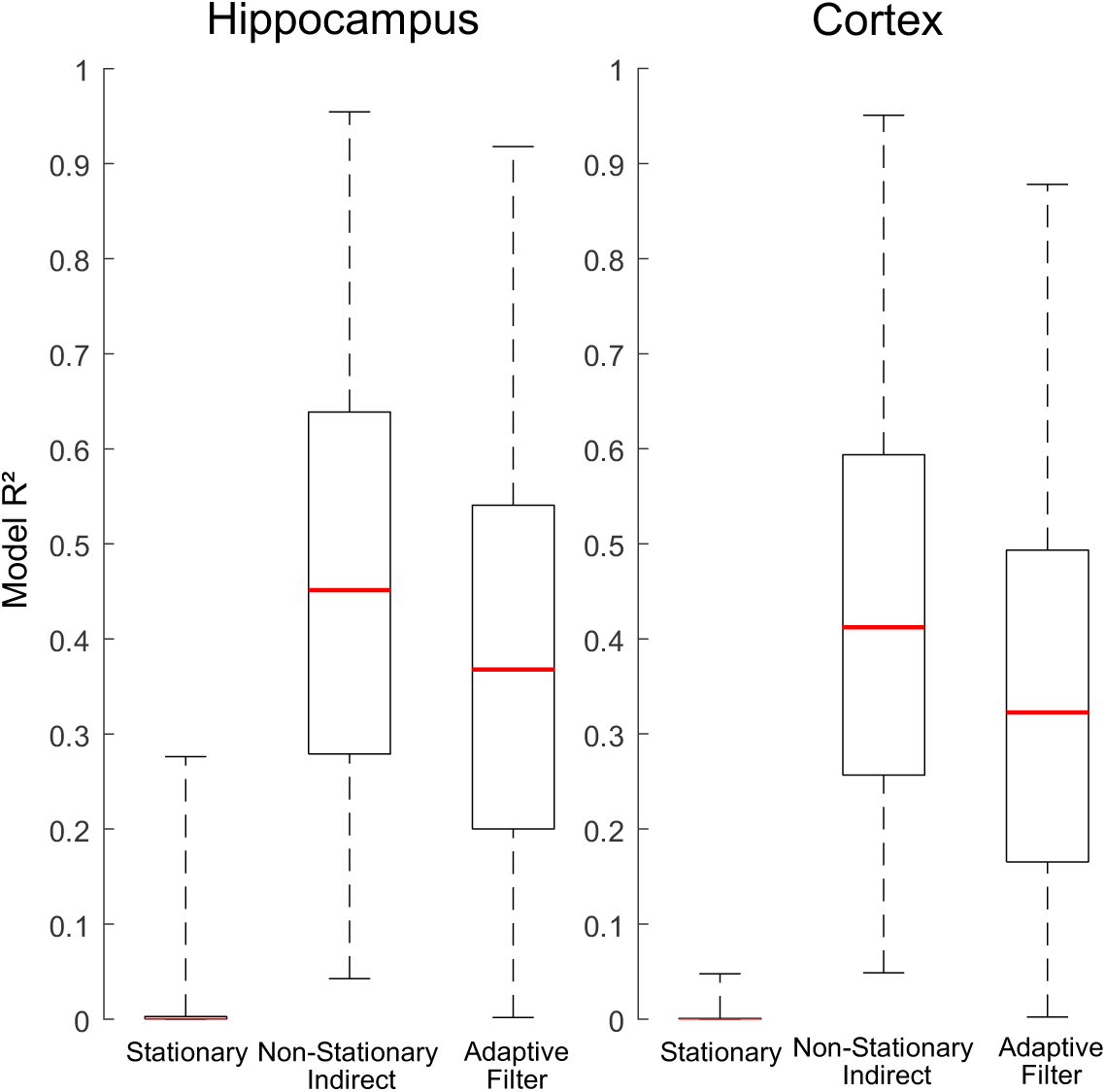
Goodness-of-fit for LFP-based models of time-varying spike-spike correlations. For hippocampus (left) and cortex (right), we fit cross-correlations measured in 60-s windows. We compare a stationary model that assumes a fixed spike-spike correlation over the recording (generalized phase, indirect optimization) to a nonstationary model that includes the local variations in LFP (generalized phase, indirect optimization), as well as an adaptive filtering model that tracks time-varying baseline and gain. Boxes denote inter-quartile ranges (i.e. middle 50% of observed model R^2^ values) and whiskers denote extrema.

## Discussion

Here we developed parametric models to predict nonstationary spike-spike correlations based on simultaneously-observed LFP. Using experimental data from mouse hippocampus and cortex, we showed how our models, with relatively few parameters, often perform as well as a previous nonparametric approach based on the spike-triggered average LFP (STA). Parametric models may also prove to be more interpretable due to the fact that they can specifically describe the contributions from distinct LFP channels and bands. We then applied these models to the problem of predicting time-varying spike-spike correlations. We found that most of the time-varying structure is due to time-varying firing rates of the neurons, and most of the structure in cross-correlogram matrices is accounted for by a rank-1 decomposition. However, there is additional variability over time that is no t well-explained by time-varying firing rates when combined with the average correlogram, and this structure can be partially-predicted by parametric models based on the LFP.

## Limitations

There are several limitations to the results presented here. Here we have modeled the influence of LFP on time-resolved spike-spike correlations without considering other factors that may also influence these correlations. Additional covariates related to stimulus or task variables could potentially be included in the parametric modeling framework used here, alongside the LFP, but it is important to note that, without including these other potentially confounding variables, the estimates of the LFP effects reflect associations rather than robust, causal estimates. Compared to nonparametric methods, the parametric methods presented here allow for greater interpretability – by which we mean there is a simple, direct relationship between specific LFP features and the spike-spike correlation. The coefficients of the parametric models directly indicate which LFP bands (e.g. alpha, theta), channels (in the two-channel model), and phases contribute to the spike-spike correlation. Here we have emphasized that in many cases, especially for the GP models, the models are also quite compact with few parameters. In the cases where more parameters are used, such as the filterbank model, it may be important to consider the possibility of overfitting. Cross-validation and, if needed, regularization penalties for the parameters (𝛽 for the indirect method, 𝛼 for the direct method) may be needed to detect overfitting and constrain these higher-dimensional models.

There are many existing statistical tools for detecting the presence of and changes in correlations (Grün, 2009; Harrison et al., 2013; Lepage et al., 2013; Zhou et al., 2015). When repeated trials are available, one common approach is to use shuffled correlations (Perkel et al., 1967; Eggermont, 1990; Chen et al., 2012), and with single trials the jittered correlogram can provide a null model. Here we have focused primarily on modeling single-trial data and changes over time with LFP-based predictors. Our parametric approach is, most directly, an alternative to nonparametric methods of linking the LFP to spike-spike correlations (Eggermont and Smith, 1995; Halliday et al., 1995; Goldberg et al., 2004; Middleton et al., 2012). However, the parametric models developed here could also be used for hypothesis testing using model comparison. When evaluating nonstationarity, it may be useful to more precisely distinguish between situations where spike-spike correlations change due to: 1) common dependence on a changing LFP, 2) changing spike-field coupling, and 3) changes unexplained by the LFP, such as plasticity in a monosynaptic connection between the neurons or due to external stimulus and task variables. The models presented here provide a straightforward approach for considering the influence of changing LFP and, with the adaptive filter, changes in the rates and shared, spike-field gain of the neurons being modeled. By extending the adaptive filter, future work could consider the possibility that the spike-field coupling of the individual neurons may itself vary over time with distinct patterns for each of the neurons. Although our results here suggest that dynamic spike-spike correlations are well described with a low-rank structure, spike-field coupling changes may be expected to occur in longer recordings and in cases where there may be more demands for adaptation and learning.

Many dynamical modeling approaches exist that could also potentially explain time-varying spike-spike correlations similar to what we have described here. GLMs that use LFP to predict the activity of single neurons (Harris et al., 2003; Canolty et al., 2010; Kelly et al., 2010) find that observed spike-spike correlations are consistent with the correlations between predicted single neuron rates (Zhou et al., 2015; Cui et al., 2016). Computing windowed correlations from the rate predictions would likely allow for nonstationary, LFP-dependent prediction of cross-correlation, as well. Additionally, state-space models and other models for shared variability, such as Poisson Linear Dynamical System (PLDS) or Gaussian Process Factor Analysis (GPFA) models (Yu et al., 2008; Li et al., 2019), may, similarly, have the capacity to describe time-varying spike-spike correlations. Although these models have not typically used LFP as a covariate, previous work has shown that they can account for overall spike-spike correlations (Macke et al., 2011) and computing windowed predictions would likely allow for nonstationarity as well. Additionally, there are several approaches from the broader problem of functional connectivity estimation (Magrans de Abril et al., 2018) that have aimed to incorporate time-varying structure, including Dynamic Conditional Correlation (DCC) methods (Engle, 2002; Lindquist et al., 2014; Hakimdavoodi and Amirmazlaghani, 2020). Although, again, these models have not typically used LFP as a covariate, they are designed to specifically track dynamic correlations. The parametric GBLM framework here, particularly the direct method, differs from these other approaches primarily in that it frames delay-resolved spike-spike cross-correlation estimates as a bilinear transformation of the covariate cross-correlations. Like the GLMs predicting single neuron activity from LFP, even a static model will show time-dependent effects due to time-varying covariates. Embedding the parametric models used here in an adaptive filtering framework provides additional flexibility similar to other models with dynamic parameters. However, adding latent variables or observed covariates beyond the LFP would likely improve the performance of the models presented here.

Another consideration in the analysis of the models here is the presence of spike bleed-through, since low-pass filtering of the wideband extracellular signal does not completely remove the effects of spikes on the LFP (David et al., 2010; Ray, 2015; Banaie Boroujeni et al., 2020). Here we have not attempted to correct for these effects, thus spike bleed-through may affect the predicted correlations for delay intervals close to 0. However, since the parametric methods are based on even-narrower filtering of the LFP, it is unlikely to play a major role in the results here, especially when *τ* ≫ 0. We have also focused exclusively on pairwise, single-trial correlations and time-domain descriptions of the spike-LFP relationship. There may be some benefits to extending our models to the population level (Yatsenko et al., 2015), using multiple trials to distinguish between spike count correlations and firing rate correlations (Vinci et al., 2016), or considering nonstationarity in the frequency domain (Zhan et al., 2006).

## Applications

Spike-spike correlations are major tools for characterizing neural processing, and many studies have documented the dependence of correlations on stimuli (Kohn and Smith, 2005), attention (Cohen and Maunsell, 2009), brain state (Ecker et al., 2014), distance between neurons, anatomical location (Senzai et al., 2019), and neuromodulation (Minces et al., 2017), among many other factors. At the same time, correlations are often used to assess potential synaptic interaction between neurons (Moore et al., 1970; Fetz et al., 1991) or functional connectivity more broadly (Gerstein and Perkel, 1969; Stevenson et al., 2008; Magrans de Abril et al., 2018). Some of these relationships, such as the effect of brain state, may be directly attributable to the LFP, while others, such as synaptic effects, are likely distinct from LFP-driven correlations. In previous work on detecting and modeling synaptic effects, researchers have explicitly attempted to distinguish between slow, time-varying baseline changes and fast synaptic effects (Ren et al., 2020; Wei and Stevenson, 2021). However, modeling the contribution of the LFP to spike-spike correlations, and how this contribution changes over time, may allow for more precise identification of both LFP-driven effects and the contributions of other factors.

Finally, one important application of the parametric models here may be in improving neural decoding. Modeling correlations improves decoding (Stevenson et al., 2012), and the presence of correlations may have a beneficial effect on readout by downstream neurons (Valente et al., 2021). Observed correlations can be also be directly used by decoders to predict stimuli (Sadeghi et al., 2019; Zhai et al., 2020), brain state (Benisty et al., 2023), and behavior (Maynard et al., 1999) more accurately than when using firing rates alone. At the same time, LFP has been used as a signal for decoding both by itself and in addition to spike signals (i.e. joint decoding). These studies have found that joint decoding generally leads to improvements, but the individual contributions from LFP vs. from spikes are partially redundant, just as the contributions from each of multiple neurons can be partially redundant. By modeling the link between LFP and observed spikes, as well as the link between LFP and spike-spike correlations, the tools developed here may allow for easier disentangling of these contributions.

## Acknowledgements

This material is based upon work supported by the National Science Foundation under Grant 1651396. Thanks to the Allen Institute for Brain Science for sharing their datasets and for supporting open science.

